# Proteostatic regulation of caveolins avoids premature oligomerisation and preserves ER homeostasis

**DOI:** 10.1101/2022.04.24.489297

**Authors:** Frederic Morales-Paytuví, Carles Ruiz-Mirapeix, Alba Fajardo, James Rae, Marta Bosch, Carlos Enrich, Brett M. Collins, Robert G. Parton, Albert Pol

## Abstract

Caveolin-1 (CAV1) and CAV3 are membrane sculpting proteins driving formation of plasma membrane caveolae. Caveola formation is unique as it requires oligomerisation of newly synthesised caveolins through the biosynthetic-secretory pathway. Here, we combine structural, biochemical, and microscopy analyses to examine the early proteostasis of caveolin family members and mutants. We describe striking trafficking differences between newly synthesised caveolins, with CAV1 rapidly exported to the Golgi but CAV3 showing ER retention and targeting to lipid droplets. Only monomeric/low oligomeric caveolins are efficiently exported from the ER, with oligomers assembling beyond the cis-Golgi and disease-causing mutations leading to detrimental non-functional complexes. Caveolins in the ER are maintained at low levels by active proteasomal degradation, avoiding premature oligomerisation and ER stress. Increasing lipid availability, cholesterol for CAV1 and fatty acids for CAV3, enhances trafficking and reduces proteasomal degradation. In conclusion, we identify proteostatic mechanisms that modulate stability and trafficking of newly synthesised caveolins, protecting cells against ER stress but perturbed in caveolin-related disease.

**Summary:** Understanding the unique proteostasis of caveolins has important implications for cell biology and physiopathology. Combining structural, microscopy, and biochemical analyses, we uncover new insights into the mechanisms that differentiate the early biosynthetic steps of caveolin family members, isoforms, and pathogenic mutants.

## Introduction

Caveolins are membrane sculpting proteins driving formation of caveolae, dynamic bulb-shaped invaginations on the plasma membrane (PM) (Parton, 2018). Caveolin-1 (CAV1) in most cell types and CAV3 in striated muscle are indispensable caveolar components. Caveolae have been linked to mechanoprotection, signalling, lipid homeostasis, and endocytosis (Cheng and Nichols, 2016). Furthermore, caveolins function independently of caveolae regulating lipid-dependent processes in the endoplasmic reticulum (ER), lipid droplets (LDs), mitochondria, and endosomes (Pol et al., 2020).

Caveola formation is unique as it requires gradual oligomerisation of newly synthesised caveolin protomers during their biosynthetic transport throughout the ER, Golgi complex (GC), and PM (Han et al., 2016). Caveolins are synthesised in the ER and, based on model systems such as *in vitro* import into microsomes, it is assumed that spontaneously oligomerise into 8S complexes (∼14 protomers), stabilised by fatty acylation or cholesterol (Han et al., 2016; Monier et al., 1996; Monier et al., 1995). The 8S complexes would be rapidly concentrated in ER-exit sites to be transported by COPII vesicles into the GC and assemble 70S oligomers (∼150 protomers), to accrue an estimated 20,000 cholesterol molecules (Hayer et al., 2010a; Ortegren et al., 2004). Finally, oligomers traffic as caveolar carriers to the PM to form long-lived but metastable caveolae (Kovtun et al., 2015; Tagawa et al., 2005).

As well as being a crucial issue related to how cells organise domains within the mosaic environment of the PM, characterisation of these biosynthetic steps has implications for understanding how disrupted proteostasis promotes pathogenesis (Hipp et al., 2019). Most naturally occurring caveolin mutations that underlie disease do so by interfering with the assembly of functional oligomers. Indeed, many caveolinopathies are defined as ER-GC disorders characterised by defective oligomerisation and formation of non-functional caveolin complexes interfering with a variety of intracellular processes such as ER-GC homeostasis, mitochondrial function, cytoskeleton organisation, and extracellular matrix formation (Gonzalez Coraspe et al., 2018). These defects lead to disease conditions such as pulmonary arterial hypertension and congenital generalised lipodystrophy in the case of CAV1 mutants or sarcolemmal damage and muscular dystrophies for CAV3 mutants (Copeland et al., 2017; Gonzalez Coraspe et al., 2018). Disrupted proteostasis could be a unifying theme explaining the variety of detrimental long-term phenotypes attributed to caveolin malfunctioning, including ageing, diabetes, or neurodegeneration (Parton et al., 2020). Hence, a full understanding of how dysfunctional caveolin biosynthetic trafficking could lead to pathogenic conditions requires a holistic characterisation of their poorly understood early proteostasis. Here, by combining structural, microscopic, and biochemical analyses to examine the initial biosynthetic steps of caveolin family members and pathogenic mutants we reveal complex proteostatic mechanisms essential for caveolin trafficking which are disrupted in disease.

## Results and Discussion

### *In silico* analysis of caveolin oligomers

While many previous studies have suggested that caveolins possess distinct functional domains, the CAV1 structure recently determined by cryoEM provides new insights into their assembly (Porta et al., 2022). Recombinant CAV1 forms a ring-shaped structure composed of 11 primarily alpha-helical subunits that spiral towards a central beta-barrel formed by their C-terminal sequence (Fig. 1A). This unique architecture produces a flat disc with one extremely hydrophobic face that likely penetrates the cytosolic leaflet of the lipid bilayer (Fig. 1B). Using the CAV1 structure as reference (Fig. 1C), we generated models of CAV1, CAV2, and CAV3 using AlphaFold2 (Evans et al., 2021; Jumper et al., 2021) and found that they are predicted to form essentially identical structures involving inter-subunit interactions (Fig. 1D, S1A and details in Methods), demonstrating the predictive capacity of the algorithm.

**Fig. 1.**
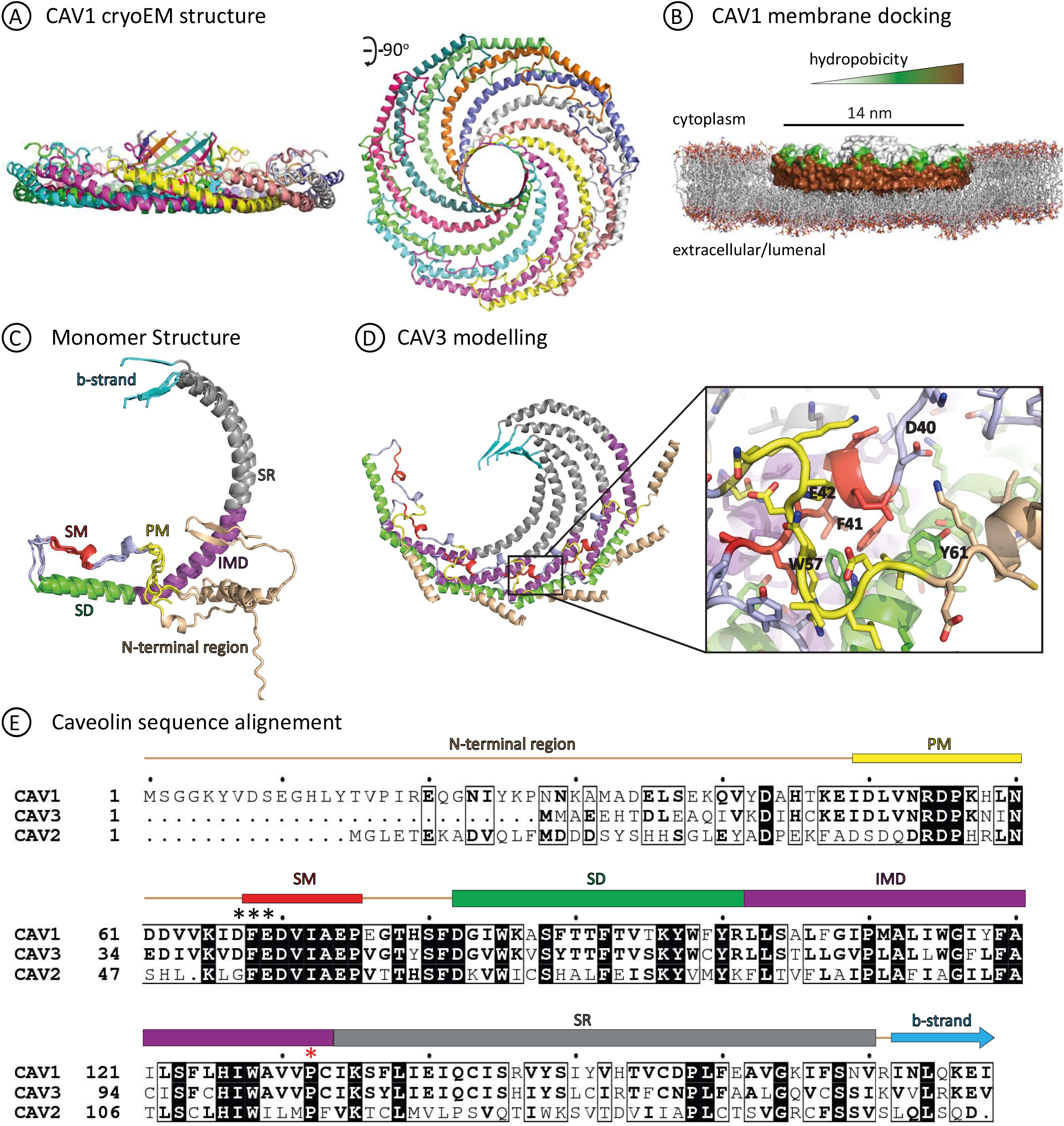
Structure of CAV1 and modelling of CAV2 and CAV3. **(A)** Structure of the CAV1 11 subunit oligomer determined by cryoEM (Porta et al., 2022). Each CAV1 protomer is shown in cartoon representation in a different colour. Note that only residues from 49-178 were resolved in the cryoEM maps. **(B)** CAV1 structure shown in surface representation coloured according to hydrophobicity. The structure was manually modelled with a lipid bilayer to show the proposed membrane docking and insertion into the cytoplasmic leaflet. **(C)** Structural alignment of individual monomers of CAV1 derived from cryoEM with models of full-length CAV1 derived from AlphaFold2 predictions. The structures are coloured using the same scheme as (Porta et al., 2022) with the positions of regions labelled: SM, signature motif (red); SD, scaffolding domain (green); and IMD, intermembrane domain (purple); PM, pin motif (yellow); SR, spoke region (grey); and beta-strand (cyan). The previously proposed oligomerisation domain (OD) contains the SM and SD. **(D)** Five subunits of full-length CAV3 modelled with AlphaFold2 form a structure essentially identical to five subunits of the CAV1 cryoEM structure (details in Fig. S1). Protomers are coloured as in panel C. The CAV3 (_40_DFE_42_) inter-subunit interactions are detailed. The central Phe (F) side chain is stacked within a hydrophobic pocket while the flanking Asp (D) and Glu (E) side chains are shielded by the PM motif of the adjacent protomer. **(E)** Sequence alignment of human CAV1, CAV2, and CAV3. Regions of the structure are indicated above and coloured as in panels C and D. The “DFE” sequence of CAV1 and CAV3 thought to mediate ER export is highlighted by asterisks (*). The site of pathogenic Pro to Leu mutation (P132L in CAV1 and P104L in CAV3) is indicated by a red asterisk.

The disc-shaped topology of tightly packed protomers provides a potential explanation for the observation that PM CAV1 is only recognised by antibodies raised against N-terminal residues – an accessible region which is not part of the ring but extends into the cytoplasm– but not by antibodies against the central signature motif or the C-terminus (Hayer et al., 2010a; Luetterforst et al., 1999; Pol et al., 2005). The latter antibodies only recognise a pool of CAV1 within the cis-Golgi complex. In contradiction to previous schemes (Han et al., 2016), these observations combined with the CAV1 structure and AlphaFold2 models (Fig. 1C and S1A) suggest that the caveolin C-terminus and signature motif remain exposed in the early Golgi, with oligomers assembled beyond this compartment.

In support of this hypothesis, the ER exit motifs previously found to act as COPII binding sequences in CAV1 (_67_DFE_69_) and CAV3 (_40_DFE_42_) (Hayer et al., 2010a) are physically inaccessible within the ring (Fig. 1, D and E). These three residues are tightly packed as part of the inter-subunit interactions required for oligomerisation. The central Phe side chain is stacked within a hydrophobic pocket while the flanking Asp and Glu side chains are shielded by the Pin motif of the adjacent protomer (Fig. 1D). Hence, oligomers formed prematurely in the ER are unlikely to be transported into the GC, suggesting *in vivo* mechanisms to avoid spontaneous oligomerisation.

### Differential biosynthetic proteostasis of caveolin family members

To test these predictions, we adapted methods previously used for characterising the early biosynthesis of caveolins (Hayer et al., 2010a) (Fig. 2A). Unlike these studies, we N-terminally tagged caveolins to avoid interfering with ring assembly or other interactions occurring in the ring centre where C-termini interact. Thus, GFP-CAV1, CAV2, or CAV3 were transfected into COS-1 cells in the presence of cycloheximide, to inhibit protein synthesis. After six hours the drug was removed to allow synchronised protein expression for three hours. We selected three hours to avoid potential artefacts only observed with longer expression times (Hayer et al., 2010a).

**Fig. 2.**
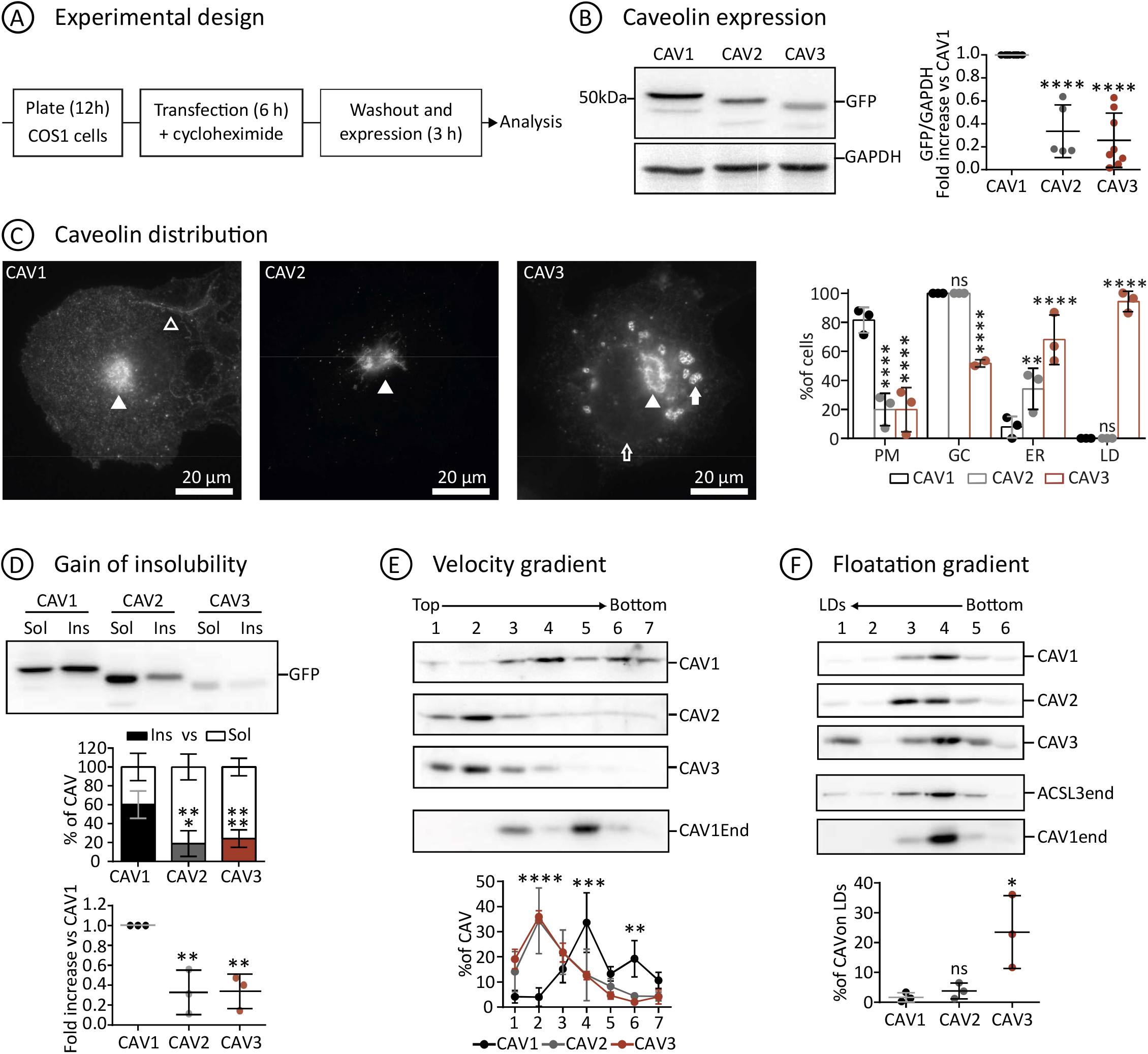
Differential proteostasis of newly synthesised caveolins. **(A)** GFP-tagged CAV1, CAV2, or CAV3 (1 μg of DNA per protein) were transfected into COS-1 cells in the presence of cycloheximide. After six hours the drug was removed to allow synchronised protein expression for three hours and cells analysed. **(B)** Caveolin levels were determined by immunoblotting (IB) of cell lysates (equal protein concentration among samples) with anti GFP antibodies. Levels were quantified by densitometry (n ≥5 independent experiments), corrected with respect to GAPDH levels in each sample, and referred to CAV1 levels. **(C)** The intracellular distribution of transfected GFP-caveolins was analysed by fluorescence microscopy. The percentage of cells in which the proteins were observed in the PM (empty arrowhead), GC (filled arrowhead), ER (empty arrow), or LDs (filled arrow) was calculated in three different experiments (n ≥ 80 cells per condition). **(D)** Transfected cells were homogenised in a buffer containing 1% TX-100 at 4ºC and soluble and insoluble fractions separated by centrifugation. Protein distribution was calculated in equal volumes of each fraction by IB with anti-GFP antibodies (n≥3 independent experiments). The upper graph shows the relative presence of caveolins in each fraction relative to the total levels of each caveolin. The lower graph shows levels of insoluble caveolin when compared with CAV1. **(E)** Transfected cells solubilized with 0.5% TX-100 were loaded at the top of sucrose density gradients and fractionated by centrifugation according to their molecular weight. Protein distribution was evaluated in equal volumes of each fraction by IB with anti-GFP antibodies (n=3 independent experiments). Sedimentation of endogenous CAV1 (CAV1end) was evaluated in non-transfected cells. Protein distribution was quantified with polyclonal antibodies in equal volumes of each fraction. **(F)** Transfected cells were homogenised in the absence of detergent, loaded at the bottom of sucrose gradients and fractionated according to their density by centrifugation. LDs floated into the top fraction (LDs). Protein distribution was calculated in equal volumes of each fraction by IB with anti-GFP antibodies (n=3 independent experiments). ACSL3 (a LD resident protein) was used to determine LD fractionation under these experimental conditions. All graphs show means ± SD; ns, not significant; *P < 0.05, **P < 0.01, ***P < 0.001, ****P < 0.0001 calculated in a one-way ANOVA [(B), (D) and (F)] or two-way ANOVA tests [(C) and (E)].

Firstly, protein expression was confirmed by immunoblotting with anti-GFP antibodies (Fig. 2B). Protein levels were significantly different, with CAV1 consistently expressed at the highest level and CAV3 at the lowest. When analysed by fluorescence microscopy, in most cells CAV1 was visible in the GC and PM (Fig. 2C). In contrast, CAV2 was largely retained in the GC. At this timepoint, CAV3 was observed in the GC but, in contrast to CAV1, mainly accumulated in the ER and LDs with rare PM labelling. The differential distribution was confirmed in C2C12 myoblasts (Fig. S2A).

Next, caveolin distribution was analysed biochemically making use of the finding that transport into the PM is associated with acquisition of insolubility to Triton X-100 (TX) (Hayer et al., 2010a). When cells were fractionated in cold detergent (1% TX at 4ºC), CAV1 demonstrated a rapid gain of insolubility, in contrast to CAV2 and CAV3 (Fig. 2D and S2A). To analyse the oligomeric state of caveolins, cells were extracted with mild 0.5% TX, loaded at the top of density gradients, and oligomers sedimented according to their molecular weight. Only CAV1 sedimented into high density fractions, reflecting oligomerisation (Fig. 2E). As expected, CAV1 (and endogenous caveolin) sedimented around two peaks, likely corresponding to 8S and 70S oligomers (Hayer et al., 2010a). In contrast, CAV2 and CAV3 remained in the top of the gradients, reflecting a monomeric or low oligomeric state. Finally, when loaded at the bottom of density gradients only CAV3 floated to the top LD-containing fraction (Fig. 2F).

Therefore, at early timepoints after synthesis, CAV1 rapidly exits the ER, oligomerises, gains insolubility, and traffics into the PM. However, although CAV2 is efficiently exported from the ER accumulates in the GC without forming oligomers. In contrast, CAV3 is retained for longer periods in the ER and stably remains in a monomeric/low oligomeric state without spontaneously forming oligomers.

### Proteasomal degradation of newly synthesised caveolins

Since CAV2 is the most divergent family member and unable to drive caveola formation alone, we focused on CAV1 and CAV3. The lower CAV3 levels when compared to CAV1 could be the result of greater exposure to degradation mechanisms in the ER (ER-associated degradation, ERAD). Active ERAD on caveolins could explain why caveolins do not spontaneously oligomerise following synthesis *in vivo*. To test if caveolins are degraded early after synthesis, CAV1 and CAV3 were expressed for three hours in the presence of the proteasomal inhibitor MG132. Inhibition led to increased caveolin levels (Fig. 3A), including CAV2 (data not shown). GFP-ALDI (an ER/LD resident hairpin protein) was also sensitive to MG132 but, at these timepoints, GFP-VAMP-Associated Protein A (transmembrane ER/GC protein) and GFP-Syntaxin 6 (GC soluble protein) were largely unaffected. Thus, in contrast to PM caveolins being long-lived proteins with an estimated half-life from 10 to 36 hours and degraded in lysosomes (Conrad et al., 1995; Hayer et al., 2010b; Ritz et al., 2011), newly synthesised caveolins have a much shorter lifetime, with up to 50% of the protein degraded by ERAD within the first three hours.

**Fig. 3.**
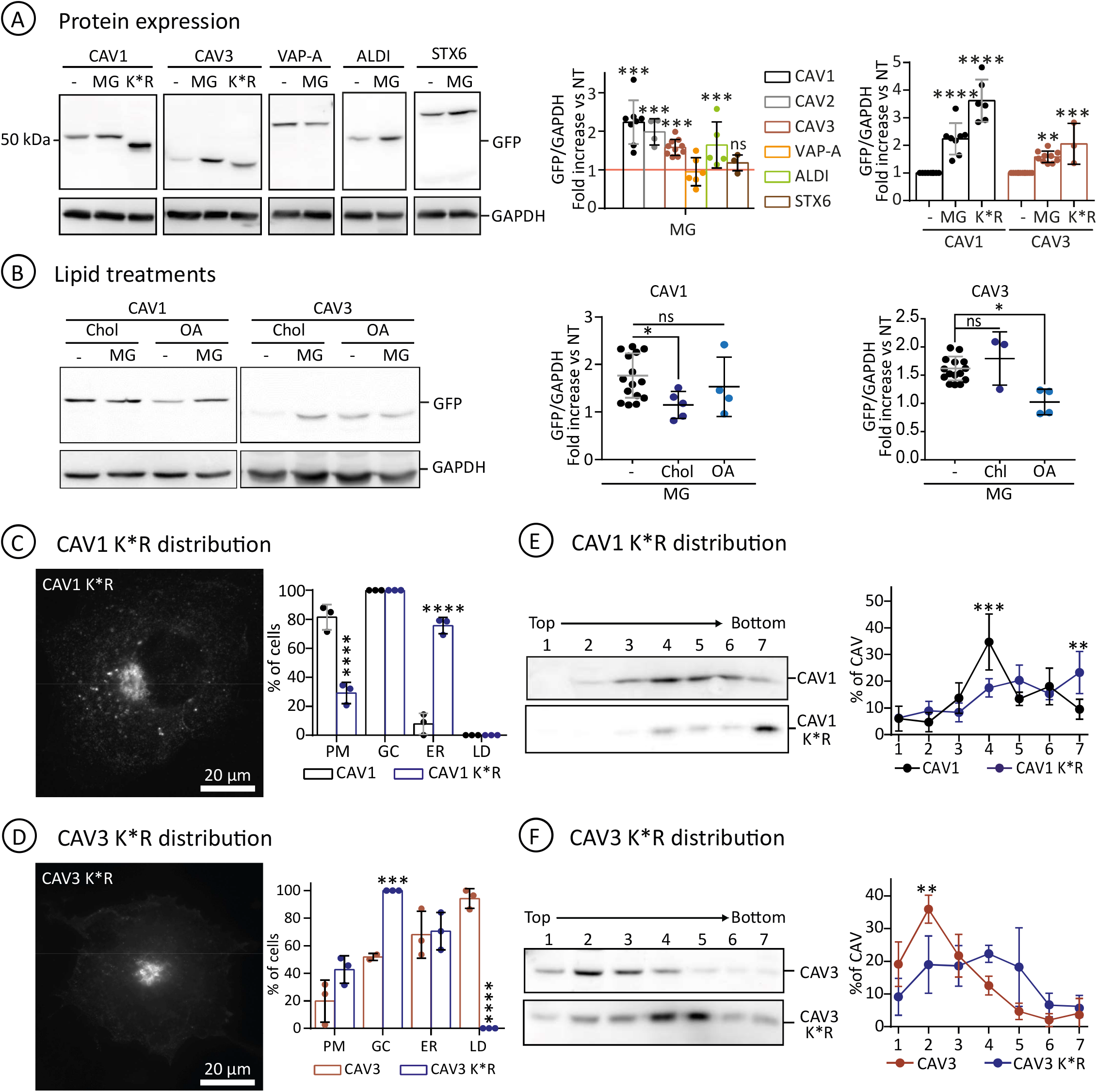
Proteasomal degradation of newly synthesised caveolin. **(A)** GFP-tagged CAV1, CAV3, VAP-A, ALDI, and STXN6 were expressed for three hours in control and MG132 treated cells. Protein levels (n≥4 independent experiments) were determined by IB with anti-GFP antibodies as in Fig. 2B and referred to the expression of each protein in the absence of MG132 (red line). CAV1 K*R and CAV3 K*R mutants were analysed in parallel (n≥3 independent experiments). All graphs show means ± SD; ns, not significant; *P < 0.05, **P < 0.01, ***P < 0.001, ****P < 0.0001 in a two-way ANOVA test. **(B)** CAV1 and CAV3 were transfected for three hours in a media additionally supplemented with cholesterol (CHL) or oleic acid (OA). Protein levels (n≥3 independent experiments) were quantified as in Fig. 2B. All graphs show means ± SD; ns, not significant; *P < 0.05, **P < 0.01, ***P < 0.001, ****P < 0.0001 in a two-way ANOVA test. **(C-F)** CAV1 K*R and CAV3 K*R were expressed for three hours, and distribution analysed as Fig. 2C in three independent experiments (n≥100 cells per condition). In addition, transfected cells were fractionated in sedimentation gradients as in Fig. 2 (n=3 independent experiments). All graphs show means ± SD; ns, not significant; *P < 0.05, **P < 0.01, ***P < 0.001, ****P < 0.0001 calculated in a one-way ANOVA test (A-B) and two-way ANOVA test (C-F).

Next, we evaluated if caveolin early proteostasis is regulated. Trafficking of CAV1 into the PM is more rapid in the presence of cholesterol and trafficking of CAV3 into LDs is proportional to fatty acid availability (Pol et al., 2004; Pol et al., 2005). Thus, we speculated that these lipids should reduce caveolin degradation by increasing its rate of trafficking out of the bulk ER. Proteasomal degradation was therefore quantified in cells supplemented with either cholesterol or with oleic acid to promote LD formation. In contrast to CAV1 in cells treated with MG132, CAV1 levels became insensitive to MG132 after cholesterol loading, suggesting that accelerated trafficking of CAV1 to the PM reduces caveolin degradation (Fig. 3B). Cells treated with oleic acid showed no significant changes in CAV1 sensitivity to MG132. In the case of CAV3, the converse was true; CAV3 degradation was unaffected by cholesterol but oleic acid addition caused loss of sensitivity to MG132 (Fig. 3B). These studies suggest differential regulation of CAV1 and CAV3 by specific lipids.

To further understand the physiological relevance of degradation of newly synthesised caveolins, and avoid the use of drugs affecting other proteins, we analysed degradation-insensitive caveolin K*R mutants (Lys substitution for Arg impairing ubiquitination). These mutants retain the capacity to traffic to the PM (Hayer et al., 2010b). When expressed for three hours, CAV1 K*R and CAV3 K*R demonstrated a higher stability than the *wt* proteins (Fig. 3, A, C, and D). When compared to CAV1, CAV1 K*R showed a reduced association with the PM while increasing its presence in the ER (Fig. 3C). The CAV3 K*R remained associated with the ER, showed increased accumulation in GC, but was excluded from LDs (Fig. 3D). In gradients, the CAV K*R mutants sedimented into heavier fractions than the *wt* proteins (Fig. 3, E and F). These results suggest that ERAD prevents premature formation of caveolin high molecular weight complexes and avoids caveolin trafficking defects.

Thus, newly synthesised caveolins are actively degraded by ERAD. Caveolin retention in the ER favours degradation and determines the net pool of caveolin trafficking into the PM or LDs. At these early time points, lipid availability inversely determines caveolin retention in the ER and the extent of degradation. In the ER, caveolins remain in a low oligomerisation state which is favoured by ERAD and required for proper trafficking.

### CAV3, but not CAV1, contains an ER-retention sequence

Only a minor pool of CAV1 is visible in the ER, likely suggesting a faster ER export when compared with CAV3 and that there may be specific mechanisms to retain CAV3. To minimise the potential sequences involved in this differential trafficking, we first confirmed that alpha- and beta-CAV1 (lacking the first N-terminal 32 residues, Fig. 1E) are identically exported from the ER (Hayer et al., 2010a). Indeed, after three hours alpha- and beta-CAV1 were indistinguishable to locate between the GC and the PM, as reflected by its insolubility, but were undetectable in the ER (Fig. 4, A, B and C). Beta-CAV1 was more stable than alpha-CAV1. Degradation of CAV1 in lysosomes is mediated by ubiquitination on Lys 5, 26, 30, 39, 47, and 57 (Kirchner et al., 2013), as beta-CAV1 lacks several of these Lys, higher stability likely reflects a less active ubiquitination and degradation.

**Fig. 4.**
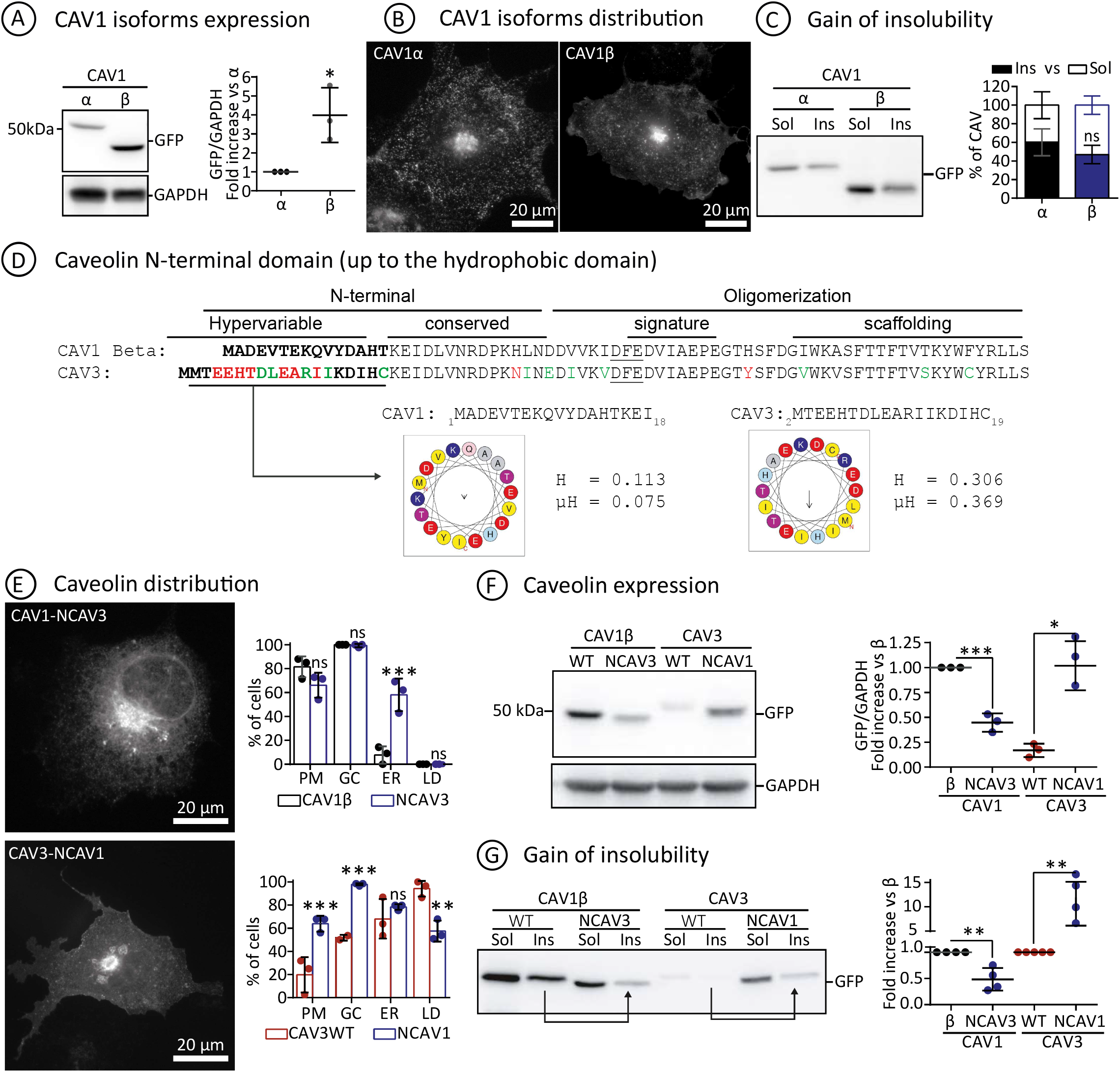
Molecular determinants for retention of CAV3 in the ER. **(A-C)** GFP-CAV1 and GFP-beta-CAV1 were expressed for three hours and protein levels analysed (n=3 independent experiments) by IB with anti-GFP antibodies (A). Protein distribution was assessed by fluorescence microscopy (B). The gain of insolubility (C) was determined as in Fig. 2. **(D)** The N-terminal domains of beta-CAV1 and CAV3, similar proteins in sequence and size, up to the hydrophobic domain (starting at LLS) are shown. Notable changes are indicated with red letters and equivalent residues with green letters. The DFE motif binding COPII is underlined. The hypervariable sequence indicates a stretch with significant differences. The hydrophobicity (H), the hydrophobic moment (μH), and tendency to fold as an alpha helix (with hydrophobic residues in yellow and charged amino acids in red and blue) of the hypervariable stretch are included. **(E)** CAV1 N-CAV3 (CAV1 with the N-terminus of CAV3) was generated by substitution of the initial 14 amino acids of beta-CAV1 by the first 16 residues of CAV3. CAV3 N-CAV1 (CAV3 with N-terminus of CAV1) was generated by substitution of the initial 16 amino acids of CAV3 by the first 14 residues of beta-CAV1. Mutants were expressed for three hours and distribution analysed in three independent experiments (n≥100 cells per condition). **(F and G)** Expression levels and gain of insolubility were analysed by IB with anti GFP antibodies as in Fig. 2 and referred to beta-CAV1 or CAV3 respectively. Arrows in G indicate the bands corresponding to the reduction or gain of insolubility of the mutants. All graphs show means ± SD; ns, not significant; *P < 0.05, **P < 0.01, ***P < 0.001, ****P < 0.0001 in a two-tailed unpaired t-test [(A) and (C)], one-way ANOVA [(F) and (G)] or two-way ANOVA tests [(E)].

Comparison of beta-CAV1 and CAV3 around the conserved COPII binding motif showed that while the signature and scaffolding domains are practically identical there is considerable divergence in the N-terminal residues (red residues, Fig. 4D). This region is enriched in hydrophobic amino acids only in the case of CAV3 and has a propensity to form an alpha-helix according to HeliQuest (Gautier et al., 2008) and AlphaFold2 (Fig. 4D, 1D and S1A). Thus, we determined if these residues retain CAV3 in the ER by exchanging the initial 14 amino acids of beta-CAV1 by the first 16 residues of CAV3 to generate CAV3 with the N-terminus of CAV1 (CAV3 N-CAV1) and CAV1 N-CAV3. When expressed for three hours, and compared with *wt* caveolins, CAV1 N-CAV3 accumulated in the GC and it was now detectable in the ER of most cells (Fig. 4E). CAV3 N-CAV1 demonstrated a lower affinity for LDs, less accumulation in GC and, in contrast to CAV3, was observed in the PM of most cells. In agreement with the redistribution, CAV1 N-CAV3 showed reduced stability, reflecting ER retention and active degradation, as well as reduced insolubility, indicating impaired arrival to the PM (Fig. 4, F and G). On the contrary, CAV3 N-CAV1 showed a higher stability and insolubility - reflecting reduced ER retention and improved trafficking to the PM.

Hence, an N-terminal alpha helix retains CAV3 in the ER. CAV1 N-CAV3 does not traffic into LDs, suggesting that in addition to ER retention other motifs mediate this lateral LD targeting. Whether the N-terminal alpha helix of CAV3 interacts with lipids or proteins in the ER deserves further studies, but these mutants confirm that ER retention reduces caveolin stability and determines the extent of protein available for the biosynthetic-secretory pathway.

### Caveolin proteostasis is maintained by ERAD and disrupted by pathogenic mutants

Our results suggest that complex proteostatic mechanisms are crucial for efficient trafficking of caveolins from and within the ER. We hypothesised that this system has evolved to maintain low levels of caveolins within the ER and minimise oligomerisation. We next investigated whether disruption of this early proteostasis is detrimental for cells. Because, in contrast to *wt* caveolins, degradation-resistant mutants prematurely form high molecular weight complexes and perturb caveolin trafficking, we determined if these defects trigger the unfolding protein response (UPR) _–_ an indicator of cellular stress. Indeed, both mutants significantly triggered the UPR, as measured by expression of the X-box binding protein 1 (XBP1) and the DNA damage-inducible transcript 3 (DDIT3 or CHOP) (Fig. 5A). The stress was more accentuated for CAV1 K*R.

**Fig. 5.**
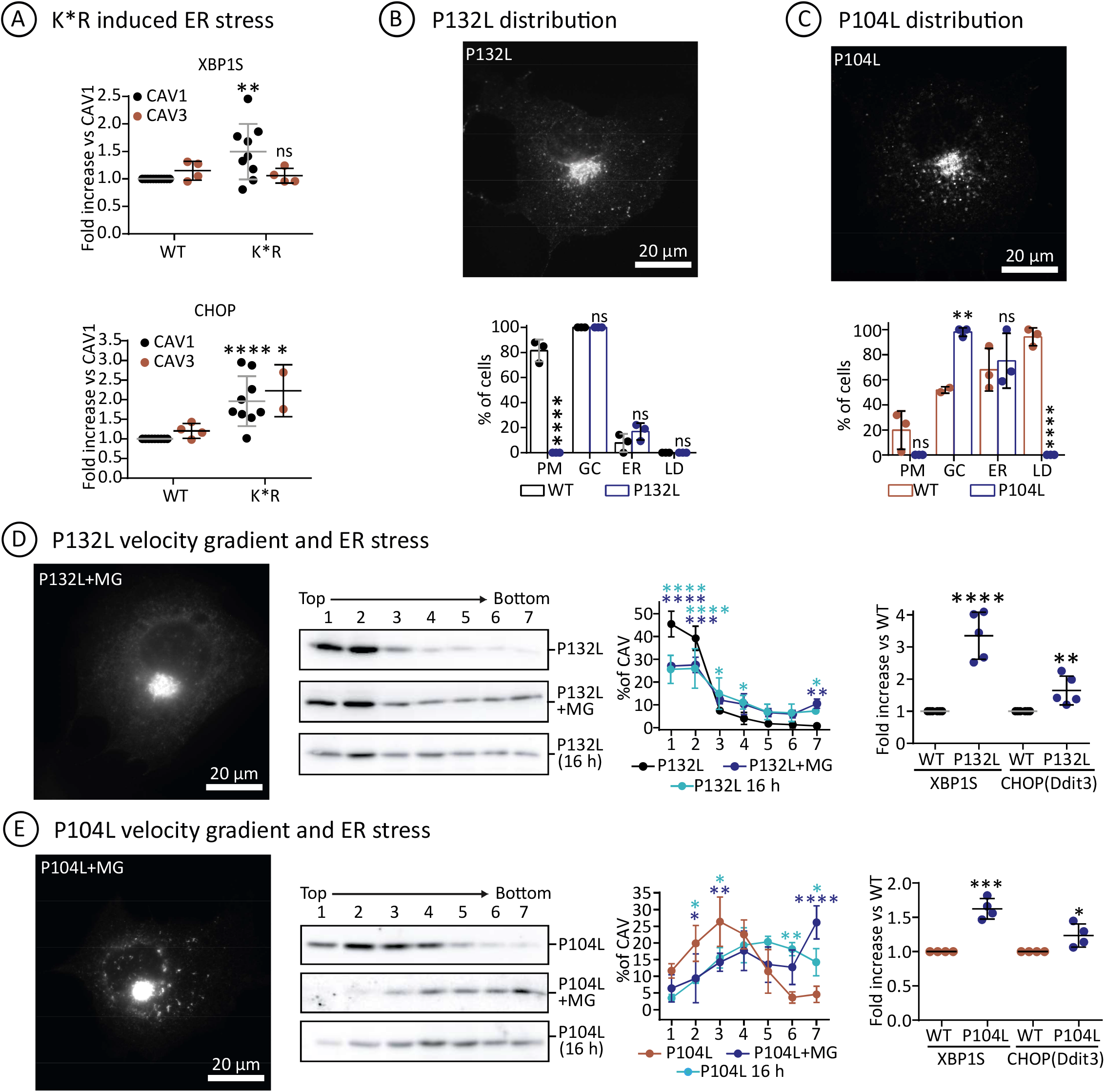
Caveolin proteostasis is disrupted by pathogenic mutants. **(A)** Wild type caveolins and CAV1 and CAV3 K*R mutants were expressed for 16 hours and the ER stress evaluated by measuring expression of XBP1 and DDIT3 (n≥4 independent experiments). Results are referred to the expression of stress markers in CAV1 transfected cells. **(B and C)** *Wt* and GFP-tagged mutants of CAV1 (P132L) and CAV3 (P104L) were transfected for three hours and distribution analysed (n=3 independent experiments including 80 cells or more per condition) (C and D) as in Fig. 2. **(D and E)** GFP-tagged mutants of CAV1 (P132L) and CAV3 (P104L) were expressed for three or 16 hours in untreated cells or for 3 hours in cells additionally treated with MG132 (n=3). A representative image of the cells treated for 3 hours with MG is included. Sedimentation of the mutants according to their molecular weight was determined as in Fig. 2. The ER stress triggered by the mutants was quantified after 16 hours expression as in Fig. 5A (n≥4 independent experiments) and referred to the respective *wt* proteins. All graphs show means ± SD; ns, not significant; *P < 0.05, **P < 0.01, ***P < 0.001, ****P < 0.0001 in a two-tailed paired t test [(A), and stress in (D) and (E)] two-way ANOVA tests [(B), (C), and distribution in (D) and (E)].

Finally, we analysed how oligomerisation defects disrupt caveolin proteostasis. Mutation of a strictly conserved Pro to Leu in the hydrophobic domain of CAV1-P132L and CAV3-P104L is a paradigmatic example (Hayashi et al., 2001; Minetti et al., 1998). Pathogenic capacity of these mutants has been commonly attributed to disruption of the assumed hairpin topology. Theoretically, substitution of a Pro, inducing a tight turn, by a Leu, forming a straight chain, would disrupt the folding needed to place both caveolin extremes facing the cytosol. However, the new structural understanding demonstrates that, rather than a hairpin, this conserved Pro plays other two other important roles. Using CAV3 as an example (Fig. S1B), the conserved Pro104 sits at the junction between the intramembrane domain and spoke region and is essential for promoting a kink in the alpha-helical structure between these two elements determining the overall caveolin curvature within the disc. Secondly, Pro104 (from chain *i*) is packed against Trp87 and Phe91 of the preceding protomer in the ring (chain *i-1*). Thus, replacing this Pro by the bulkier beta-branched side chain of Leu likely perturbs both curvature and inter-subunit packing.

The early proteostasis of GFP-CAV1-P132L and GFP-CAV3-P104L was analysed after 3 hours. Like CAV1, CAV1-P132L was rarely visible in the ER and accumulated in GC and thus, oligomerisation is dispensable for this transport. However, CAV1-P132L was completely undetectable on the PM and accordingly, oligomerisation is mandatory in this step (Fig. 5B). Like CAV3, CAV3-P104L distributed between the ER and GC but in contrast to CAV3, it was absent in the PM and rarely visible on LDs (Fig. 5C). Proteasomal inhibition made evident the ER pool of both mutants, indicating active ERAD. Although accumulated in the GC, both mutants demonstrated impaired oligomerisation (Fig. 5, D and E). Proteasomal inhibition, or longer expression times (16 hours), caused progressive formation of high molecular weight complexes (Fig. 5, D and E). Both mutants triggered the UPR and similarly to K*R mutants, stress was more pronounced for CAV1-P132L (Fig. 5, B and C). After 16 hours, CAV1-P132L largely remained in low molecular weight species (Fig. 5D), suggesting that accumulation of caveolin monomers in GC also has detrimental effects.

Therefore, CAV1-P132L and CAV3-P104L are efficiently exported as monomers or lower order oligomers from the ER into the GC. In the GC the Pro to Leu substitution impairs formation of functional oligomers likely by disrupting curvature and inter-subunit packing. These mutants gradually aggregate into GC-retained and non-functional high molecular weight complexes and progressively trigger ER stress, with a likely contribution of caveolin monomers.

### Concluding Remarks

Our model cell system of synchronised caveolin expression followed by analyses at very early time points has identified several novel proteostasis mechanisms differentially regulating caveolin family members, isoforms, and mutants.

First, in contrast to the current view, we propose that caveolins are transported from the ER into the GC as monomers or small oligomeric species and 11-mer (8S) rings are only assembled beyond cis-Golgi. These dynamics are (i) predicted by structural analysis showing that residues binding COPII are buried and inaccessible once rings are formed and thus, oligomers are likely retained in the ER; (ii) illustrated by our previous observations that antibodies against the signature motif or C-terminal residues - which are hidden within oligomers – still recognise the Cis-Golgi pool of caveolin but not oligomers in the PM (Luetterforst et al., 1999; Pol et al., 2005); and (iii) exemplified by Pro to Leu mutants that - although failing to oligomerise-are exported from the ER into the GC as efficiently as *wt* caveolins. These mutants remain as monomers because the bulkier beta-branched side chain of Leu perturbs a hydrophobic pocket required for interacting with neighbouring protomers. Accumulation of caveolins in the ER and GC results in the gradual formation of non-functional aggregates and causes stress, a detrimental process also promoted by the accumulation of monomers.

Second, in support of the above conclusion and exemplified by CAV3, we describe that caveolins residing in the ER remain in a low oligomeric state. This is possible because the spontaneous oligomerisation observed *in vitro* is avoided *in vivo* by active ERAD of newly synthesised caveolins. ERAD is functioning on monomers, as illustrated by CAV1-P132L, and potentially on oligomers because most ubiquitinated Lys of caveolin remain exposed outside ring. Targeted mutagenesis to inhibit degradation causes premature formation of high molecular weight species, trafficking defects and ER stress. Although its role is primarily to eliminate detrimental misfolded proteins, in lower eukaryotic systems ERAD additionally determines distribution of normally folded proteins among different destinations (Ruggiano et al., 2016). Our studies suggest that ERAD regulates caveolin trafficking throughout the biosynthetic-secretory pathway and into LDs.

Third, we demonstrate significant differences in the early trafficking of CAV1 and CAV3; with CAV1 rapidly transported into the PM but CAV3 retained in the ER by an N-terminal alpha helix for longer periods. Cholesterol availability regulates CAV1 stability after synthesis by accelerating its exit from the ER and reducing ERAD to increase the post-GC pool. CAV1 is transcriptionally regulated by cholesterol (Hailstones et al., 1998), directly binds cholesterol (Murata et al., 1995), and caveolin deficient cells accumulate cholesterol in the ER and GC (Bosch et al., 2011a; Bosch et al., 2011b). We speculate that the stability of CAV1 could function as a cholesterol sensor, mediating cholesterol transport to the PM and, once in caveolae, adjusting the pool of accessible cholesterol for the rest of the PM (Pol et al., 2020).

In contrast, the stability of newly synthesized CAV3 responds to the presence of fatty acids that increase the LD pool and reduce its degradation. The CAV3 gene promoters are regulated by the retinoic acid receptor (RAR)-related orphan receptor alpha (RORα) (Lau et al., 2004), a transcription factor involved in fatty acid metabolism, activation of mitochondria, and regulation of genes involved in circadian rhythms. Thus, the high affinity of newly synthesised CAV3 for LDs could be part of a bioenergetic program, designed specifically for muscles, to rapidly respond to fluctuations in fatty acid availability occurring, for example, during circadian cycles, starvation, or exercise.

In conclusion, these studies emphasise the essential role of proteostatic mechanisms in the life cycle of caveolins and identify key differences between family members. Perturbation of these mechanisms allows precocious oligomerisation causing ER stress. This explains the well-documented harmful effects of high expression of *wt* caveolin (Hanson et al., 2013). Interestingly, specialised mechanisms also exist for specific degradation of mature caveolin oligomers in post-Golgi endosomal compartments (Hayer et al., 2010b; Ritz et al., 2011), emphasising the precise regulatory mechanisms that have evolved to control caveolin levels at various stages of its trafficking itinerary, both before and after reaching the PM. The identification of these mechanisms provides new insights into the effect of caveolin pathogenic mutants which show dysregulation of the sequential biosynthetic process required for assembly of caveolin oligomeric rings in the GC and formation of mature caveolae. Disrupted caveolin proteostasis, either directly caused by mutations in caveolins or indirectly when normal caveolins are affected by other environmental stresses, could be a unifying theme explaining the variety of detrimental long-term phenotypes attributed to caveolin dysfunction; including ageing, diabetes, or neurodegeneration.

## Materials and Methods

### Plasmids

pEGFP-C3-CAV1m, pEGFP-C1-CAV3m, pEGFP-C1-CAV1m P132L and pEGFP-C1-CAV3m P104L were purchased from Genscript (Piscataway, New Jersey). pEGFP-C1-CAV2m was obtained by subcloning pCMV6-Myc-DDK-CAV2m (RC202703, Origene, Rockvylle, Maryland) into a pEGFP-C vector using primers containing BglII and HindIII sites. mEGFP-CAV1h K*R was obtained from Addgene (27766, Watertown, Massachussets). pEGFP-C1-CAV3m K*RT was derived from the plasmid pUC57-CAV3m K*R acquired from Genscript, and subcloned into PEGFP-C1 vector using BglII and HindIII sites. pEGFP-C3-CAV1mβ starting at MADEVT was subcloned from pEGFP-C3-CAV1m using primers containing BsrgI and BamHI sites. peGFP-C3-CAV1m-NCAV3m was designed by replacing the hypervariable region of CAV1m β (MADEVTEKQVYDAHT), with the N-terminal region of CAV3 (MMTEEHTDLEARIIKDIHC). Both sites were swapped after a Bsrgl/BamHI digestion. pEGFP-C1-CAV3m-NCAV1m β was obtained by replacing the hypervariable region of CAV3 (MMTEEHTDLEARIIKDIHC), with the N-terminal region of CAV1mβ (MADEVTEKQVYDAHT). Both sites were swapped after a BsrglI/BamHI digestion.

### Protein structural prediction, modelling and visualization

The structural predictions of human caveolin paralogues were performed using the AlphaFold2 neural-network (Jumper et al., 2021), implemented within the ColabFold pipeline (Mirdita et al., 2021), using default settings, and using MMseqs2 for multiple sequence alignments (Mirdita et al., 2019). Sequence accession numbers are provided in **Table S1**. Monomeric predictions were performed using both AlphaFold2-ptm and AlphaFold2-multimer-v2 and compared to cryoEM structure of CAV1 monomers. AlphaFold2-ptm was used for the modelling of larger homooligomers with varying number of subunits and disc-like structures for all the caveolins. Homopentameric CAV1, CAV2, and CAV3 were focused due to the high similarity of CAV1 cryoEM and their inter-subunit interactions were analysed from i-2 to i+2 ring. Surface hydrophobicity images were mapped using Protein-Sol Patches server (Hebditch and Warwicker, 2019) and structures were rendered with Pymol (Schrodinger, USA; https://pymol.org/2/).

### Cells and reagents

COS1 (ATCC CRL-1650) and C2C12 (ATCC CRL-1772) were cultured in Dulbecco’s modified Eagle’s medium (DMEM, Biological industries, Cromwell, Connecticut) supplemented with 10% v/v foetal bovine serum (Biological industries) and 4mM L-glutamine, 1mM pyruvate (Sigma-Aldrich, St Louis, Missouri), 50 U/mL penicillin, 50 μg/mL streptomycin and non-essential amino-acids (Biological industries). Oleic acid (O1008, Sigma-Aldrich) was conjugated with fatty-acid free BSA (A8806, Sigma-Aldrich) at a 6:1 molar ratio. Cholesterol (C8667, Sigma-Aldrich) was conjugated with Methyl-β-Cyclodextrin (C8667, Sigma-Aldrich) at a 1:20 molar ratio. Cycloheximide (CHX, C7698) was purchased from Sigma-Aldrich. MG132 (MG) was purchased from Merck (474790, Kenilworth, Nova Jersey).

### Cell transfection and treatment

GenJet™ Plus (Signagen, Frederick, Maryland) and Lipofectamine® LTX (Invitrogen, Carlsbad, California) reagents were used to transfect COS1 and C2C12 cells respectively following the manufacturer’s instructions. Before the addition of the lipid-DNA complex, 50 μg/mL CHX were added. After 6 h of incubation, washout of CHX was performed by three washes with complete media and gently tapping. Synchronised protein expression pulse was expressed with or without 50 μM MG132, 1.1 mM OA or 4 mM cholesterol and chased at 3h

### Total cell lysates

Transfected cells were washed twice with cold PBS and gently scrapped with 80 µL of lysis buffer (50 mM Tris HCl pH 7.5, 150 mM NaCl and 5 mM EDTA and protease and phosphatase inhibitors) with 0.1% Triton X-100 (T8787, Sigma-Aldrich). Collected samples were completely disrupted by sonication for 20 s at 30 Amps and protein concentration was quantified using the Bio-Rad protein assay kit (500-0006, Bio-Rad, Hercules, California). 30-60 μg of protein were loaded on an SDS-page gel.

### Triton solubilization assay

The protocol was adapted from (Schlegel et al., 1999) and performed on ice. Transfected cells were washed two times with chilled PBS and solubilized for 35 min without agitation with 300 μL of cooled lysis buffer with 1% Triton X-100. Soluble fraction was collected by decantation and pipetting. Insoluble fraction was collected by scrapping cells with 300 μL of SDS lysis buffer (2% SDS, 1%Tris-HCl pH6.4) and further disrupted by passing the lysate through a 25 G needle 10 times. 30 μL of all fractions were loaded on an SDS-page gel.

### Sucrose velocity gradient

The protocol was adapted from (Hayer et al., 2010a). 1×10^6^ cells grown in a 60 mm well plates were washed two times with PBS, gently scrapped in 500 µL PBS and centrifuged at 800 G for 4 min at RT. Cells were permeabilized in 200 μL of lysis buffer 0.5% Triton X-100 for 20 min and centrifuged for 5 min at 1,100 G to discard insolubilized cells. 200 μL of supernatant were loaded in the upper phase of a sucrose step gradient of 500 μL of 40%, 30%, 20% and 10% sucrose in lysis buffer, in a polypropylene centrifuge tube (347357, Beckman Coulter, Brea, California). Gradient tubes were then ultracentrifuged (Sorval MX 150, Thermo Scientific) at 55,000 rpm for 4 h 15 min at 4ºC (Hitachi rotor, 55S, Tokyo, Japan). Seven fractions of 310 µL were collected from top to bottom and 30 μL of all fractions were loaded on an SDS-page gel.

### Sucrose flotation gradient

Two 100mm plates with 3×10^6^ transfected cells treated with 1.1 mM Oleic Acid for 3 h after CHX washout were used for each condition. Cell’s lysates were obtained by gently scraping in 300 μL of lysis buffer followed by cavitation at 55 Bar for 15 min at 4ºC. Samples were passed throughout a 22 G needle 25 times and centrifuged at 1500 G for 5 min at 4ºC. 500 µL of the supernatant fraction were mixed with 500 µL of 65% sucrose and place at the bottom of a step gradient of 200 μL of 30%, 25%, 20%, 15%, 10%, 5% (w/v) sucrose in lysis buffer and ultracentrifuged (Sorval MX 150, Thermo Scientific) at 50,000 rpm for 3 h at 4ºC (Hitachi rotor, S55S). Five fractions of 300 µL were collected using a tube slicer from top to bottom and 700 μL were collected from the bottom phase. 30 μL of all fractions were loaded on an SDS-PAGE gel.

### SDS-PAGE and Immunoblotting

Protein samples were loaded onto a polyacrylamide gel using a Bio-rad Mini-Protean II electrophoresis cell and transferred to a nitrocellulose membrane. Membranes were blocked in 5% non-fat dry milk in TBS-T for 1 h and incubated with the primary antibodies anti-GFP (1:5,000; ab290, Abcam, Cambridge, UK), anti-GAPDH (1:5,000; A00191, Genescript), and anti-CAV1 (1/3,000; BD Biosciences, San José, California). Membranes were washed and incubated with peroxidase-conjugated secondary antibodies (1:3,000; Bio-Rad) and detected with ECL (Biological Industries). Western blot quantification was performed using the Image J software (NIH).

### Fluorescence microscopy

Cells were grown in coverslips and fixed for 1h with 4% PFA and mounted with Mowiol mounting media (475904, Calbiochem, La Jolla, San Diego). Images were taken with Leica AF600 microscope (Leica Microsystems, Manheim, Germany) or Zeiss Axiovert 200 UprightMicroscope Stand (Zeiss, Oberkochen, Germany) with LSM 710 Meta Confocal Scanner using the x63 oil immersion objective lens. Images were analysed using the Adobe Photoshop CS3 software (Adobe Systems Inc, San Jose, California) and ImageJ (1.52i, National Institutes of Health, USA).

### Gene expression by quantitative PCR (qPCR)

Cells in 35mm well plates were washed two times with PBS and RNA was isolated using the EZ-10 DNAway RNA Mini-Preps kit (BS88136, Biobasic, Toronto, Canada) according to manufacturing instructions. cDNA was synthetized from 1μg RNA using the High-Capacity cDNA Reverse Transcription Kit (Applied Bioscience, ThermoFisher Scientific). qRT-PCR was performed using SYBRGreen-based technology GoTaq® qPCR master-mix (A600A, Promega, Madison, Wisconsin) and LightCycler 96® Real-time PCR system (Roche, Basilea, Switzerland).

### Statistical analysis

All data shown in graphs are the mean ± SD. Statistical significance was determined using paired t test, one-way and two-way analysis of variance (ANOVA) multiple comparisons test as specified in Fig. legends [not significant (ns), *P < 0.05, **P < 0.01, ***P < 0.001, and ****P < 0.0001]. The number of independent experiments (n) is indicated in each Fig. legend.

### Fig. preparation

Fig. were created using Microsoft PowerPoint (Microsoft 365MSO) and Adobe Illustrator (Adobe systems, v25.1.0.90). Images were edited with Adobe Photoshop CS3 software (Adobe Systems) and Fiji/ImageJ (1.52i, National Institutes of Health, USA). GraphPad Prism 7 (GraphPad Software) was used to create graphs and calculate statistical significances.

## Acknowledgements

AP was supported by the Ministerio de Ciencia e Innovación (MICINN, RTI2018-098593-B-I00) and the CERCA Programme/Generalitat de Catalunya. R.G.P. is supported by the National Health and Medical Research Council (NHMRC) of Australia (program grant, APP1037320 and Senior Principal Research Fellowship, 569452). B.M.C. is supported by an NHMRC Senior Research Fellowship (APP1136021) and an ARC Discovery Project Grant (APP1181135). FM-P is supported by the FPI program (MINECO, BFU2015-66401-R). We thank Dr. Dominic Hunter and the members of the Parton, Collins, and Pol groups for critical reading of the manuscript. We would like to thank Melanie Ohi and Anne Kenworthy for providing the CAV1 structural coordinates prior to public release. We would also like to acknowledge and thank Milot Mirdita, Martin Steinegger and the ColabFold team for making their pipeline available for public use and Drs. Vikas Tillu and Michael Healy for discussion of structural modelling.

## Author Contribution

Conceptualization: R.G.P., and A.P. Methodology: F.M-P., C.R., M.B., B.M.C. and A.F. Formal analysis: M.B., R.G.P., B.M.C. and A.P. Investigation: F.M-P., A.F., J.R., and C.R-M. Resources and supervision: R.G.P., and A.P. Data curation: F.M-P., M.B., R.G.P., and A.P. Writing (original draft): A.P. Writing (review and editing): R.G.P., B.M.C. and A.P. Visualization: B.M.C., R.G.P. and A.P. Funding acquisition: R.G.P., and A.P.

## Legends to Supplementary Figures

**Fig. S1.**
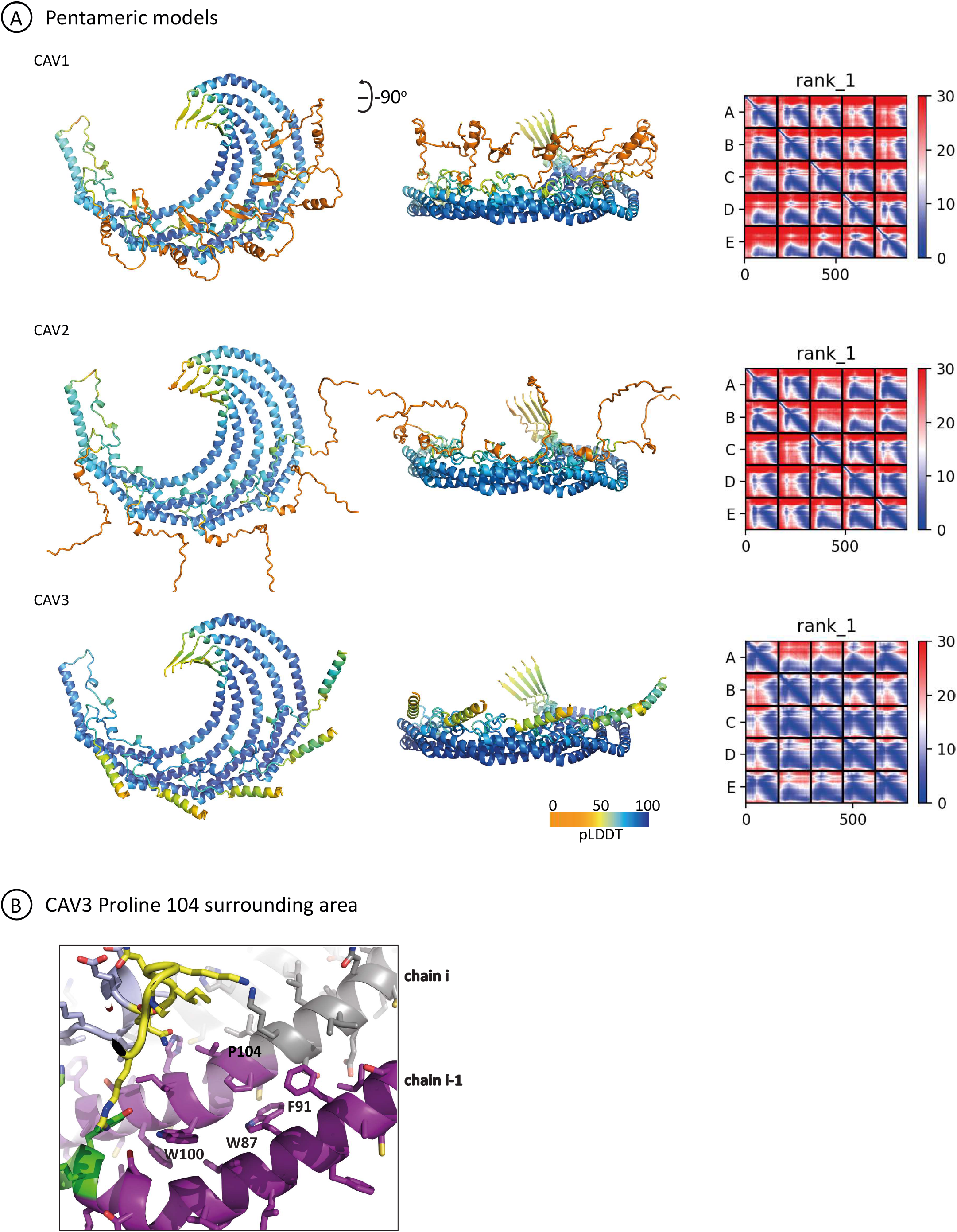
AlphaFold2 predictions of pentameric homooligomers of human CAV1, CAV2 and CAV3. **(A)** The top-ranking structures of homo-pentameric models of human CAV1, CAV2, and CAV3 predicted by AlphaFold2 are shown in ribbon diagram and coloured according to the pLDDT scores. The right-hand panels show the plots of the Predicted Alignment Error (PAE) for each top-ranking model. There is a strong degree of correlation between the five chains in each structure indicating their physical association with each other. **(B)** Close up of the region surrounding Pro105 in CAV3 in the predicted oligomeric state.

**Fig. S2.**
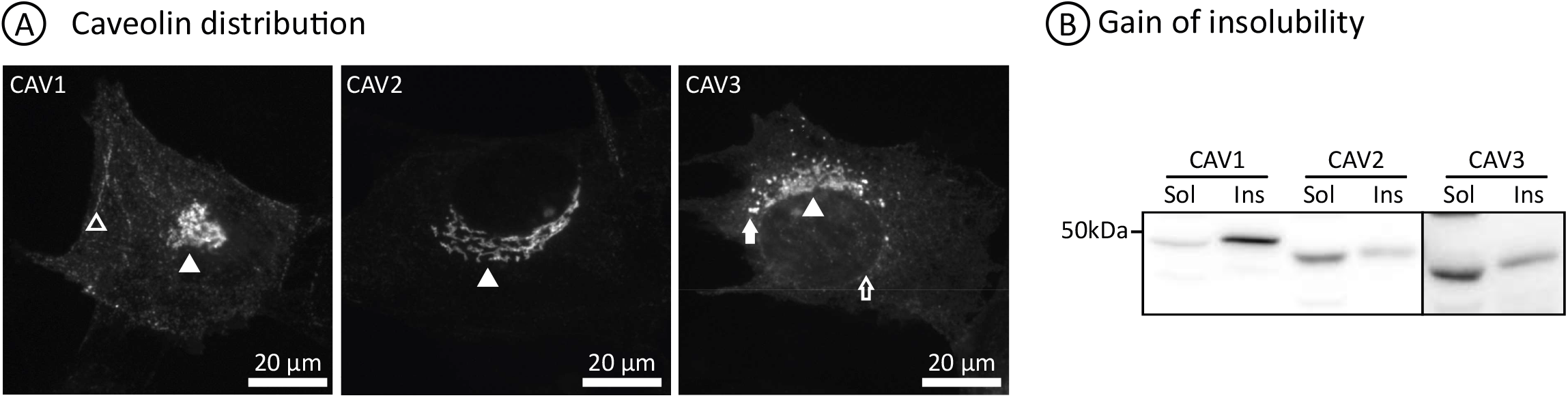
CAV1 distribution. GFP-tagged CAV1, CAV2, and CAV3 were expressed for three hours in C2C12 muscular cells. (**A**) Representative images of fluorescence staining are shown with the plasma membrane (PM), Golgi Complex (GC), Endoplasmic Reticulum (ER), and Lipid Droplets (LDs) indicated with open arrowhead, arrowhead, open arrow and arrow respectively. (**B**) Representative images of gain of insolubility are shown.

**Table S1.**
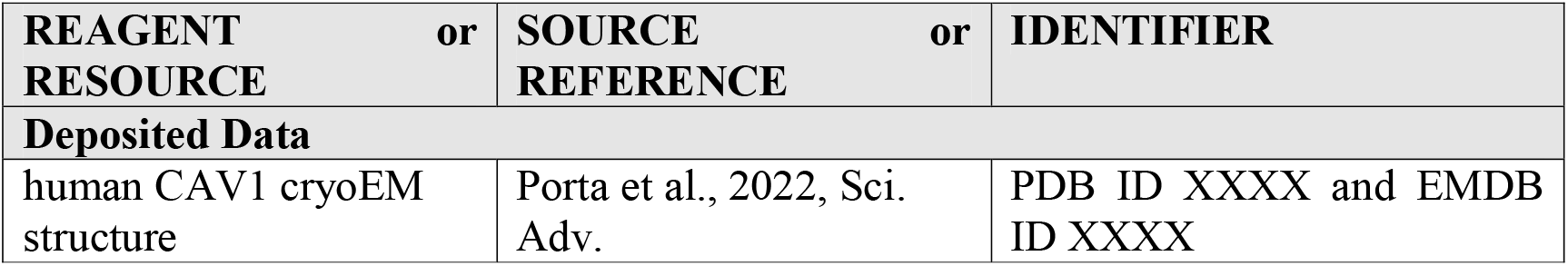
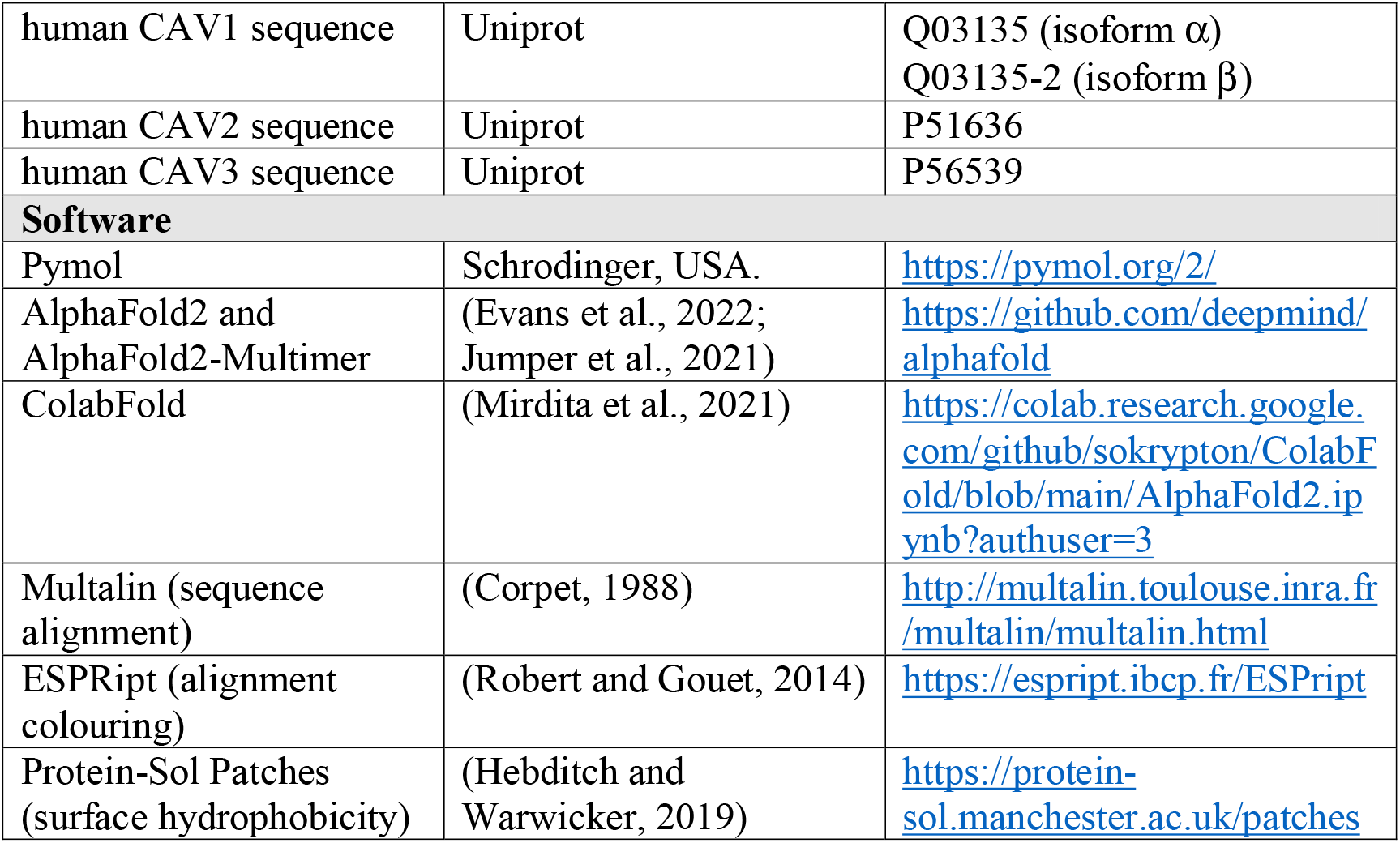
Key Resources Table.

